# Initial-state-dependent, robust, transient neural dynamics encode conscious visual perception

**DOI:** 10.1101/133983

**Authors:** Alexis T. Baria, Brian Maniscalco, Biyu J. He

## Abstract

Recent research has identified late-latency, long-lasting neural activity as a robust correlate of conscious perception. Yet, the dynamical nature of this activity is poorly understood, and the mechanisms governing its presence or absence and the associated conscious perception remain elusive. We applied dynamic-pattern analysis to whole-brain slow cortical dynamics recorded by magnetoencephalography (MEG) in human subjects performing a threshold-level visual perception task. Up to 1 second before stimulus onset, brain activity pattern across widespread cortices significantly predicted whether a threshold-level visual stimulus was later consciously perceived. This initial state of brain activity interacts nonlinearly with stimulus input to shape the evolving cortical activity trajectory, with seen and unseen trials following well separated trajectories. Furthermore, cortical activity trajectories during conscious perception are fast evolving and robust to small variations in the initial state. These results suggest that brain dynamics underlying conscious visual perception belongs to the class of initial-state-dependent, robust, transient neural dynamics.

**AUTHOR SUMMARY:** What brain mechanisms underlie conscious perception? A commonly adopted paradigm for studying this question is to present human subjects with threshold-level stimuli. When shown repeatedly, the same stimulus is sometimes consciously perceived, sometimes not. Using magnetoencephalography, we shed light on the neural mechanisms governing whether the stimulus is consciously perceived in a given trial. We observed that depending on the initial brain state defined by widespread activity pattern in the slow cortical potential (<5 Hz) range, a physically identical, brief (30 - 60 ms) stimulus input triggers distinct sequences of activity pattern evolution over time that correspond to either consciously perceiving the stimulus or not. Such activity pattern evolution forms a “trajectory” in the state space and affords significant single-trial decoding of perceptual outcome from 1 sec before to 3 sec after stimulus onset. While previous theories on conscious perception have emphasized sustained, high-level activity, we found that brain dynamics underlying conscious perception exhibit fast-changing activity patterns. These results significantly further our understanding on the neural mechanisms governing conscious access of a stimulus and the dynamical nature of neural activity underlying conscious perception.

## INTRODUCTION

Identical sensory input can be processed by the brain consciously or unconsciously [1, 2]. What determines whether a sensory stimulus gains conscious access or not? And what distinguishes neural activity underlying conscious and unconscious processing? Traditionally, neuroscientists have studied these two questions separately, by investigating how pre-stimulus activity and stimulus-evoked activity (i.e., changes in post-stimulus activity from the pre-stimulus baseline) correlate with the state of perception, respectively. However, these questions remain unsatisfactorily addressed, and a unified framework that accounts for both pre- and post- stimulus findings in an integrated manner is currently lacking.

In the pre-stimulus period, prior studies have shown that pre-stimulus amplitude of alpha and gamma-band activity [3, 4] and pre-stimulus phase of alpha oscillations [5, 6] in posterior visual regions influence perceptual outcome during threshold visual perception, suggesting that fluctuating excitability of sensory areas influences conscious access. Recent evidence also suggests that the phase of slow cortical potentials (SCPs, <5 Hz) in widely distributed cortical areas at stimulus onset influences conscious access of a threshold-level stimulus [7, 8]. However, the mechanism of this latter observation – how exactly these pre-stimulus activity patterns influence post-stimulus processing and, consequently, conscious perception – remains elusive. Additionally, whether pre-stimulus brain activity influences unconscious processing of a stimulus remains largely unexplored.

In the post-stimulus period, previous studies have reported that late-onset (> 200 ms), long-lasting (up to ~1 sec) neural activity using a variety of measures – event-related field/potential [7, 9–11], power of gamma-frequency range [11–13], and single-neuron firing [14, 15] – correlates with conscious visual perception. These findings have led to the suggestion that sustained, high-level activity across widely distributed brain regions underlies conscious processing [16, 17]. However, whether this long-lasting activity is sustained and steady or, alternatively, fast changing remains unclear [18, 19]. Furthermore, how pre-stimulus activity contributes to the presence or absence of this neural signature remains unknown.

Here we applied a dynamical systems framework [20–27] to whole-head MEG signals recorded from human subjects performing a threshold visual perception task to identify distinguishing features between conscious and unconscious processing. We characterize the dynamics of widely distributed neural activity by studying the evolution of its trajectory through multi-dimensional state space and the encoding of perceptual outcome in single-trial activity. We observed that before stimulus onset, large-scale neural activity in the SCP range was well separable at the single-trial level between seen and unseen conditions. Depending on this initial state, stimulus input triggers large-scale SCP activity to follow distinct trajectories (corresponding to different sequences of spatial patterns) in seen and unseen trials, with the activity pattern evolving far more quickly in seen trials. Importantly, seen and unseen trials were most discriminable when the population activity was changing the fastest, suggesting that transient dynamics [23, 26, 28] instead of steady-state activity characterizes these neural dynamics. In addition, transient neural dynamics during conscious perception exhibited strong across-trial variability reduction, suggesting that they are robust to small variations in the initial state. By contrast, large-scale neural activity in higher-frequency ranges did not distinguish between seen and unseen trials. Together, these results suggest a parsimonious framework that explains pre-stimulus and post-stimulus activity differentiating conscious from unconscious processing in an integrated manner, embodied in initial-state-dependent, robust, transient dynamics in the SCP range. Last but not least, we found that pre-stimulus slow cortical dynamics also influences unconscious perceptual decision making about a fine-grained stimulus feature.

## RESULTS

On average, subjects reported seeing the stimulus in 48.9 ± 4.4% (mean ± s.e.m.) of trials, and answered correctly about its orientation in 79.5 ± 2.5% of trials (Fig. 1B). When they reported seeing the stimulus, they correctly identified its orientation in almost all (96.8 ± 1.3%) of the trials. Given the small number of seen yet incorrect trials, these were excluded from all analyses. When they denied seeing the stimulus, they guessed correctly about its orientation in 62.0 ± 2.7% of trials, which was still significantly above chance level at 50% (*p* = 0.001, Wilcoxon signed rank test), indicating unconscious processing of the stimulus [29, 30].

**Fig. 1.**
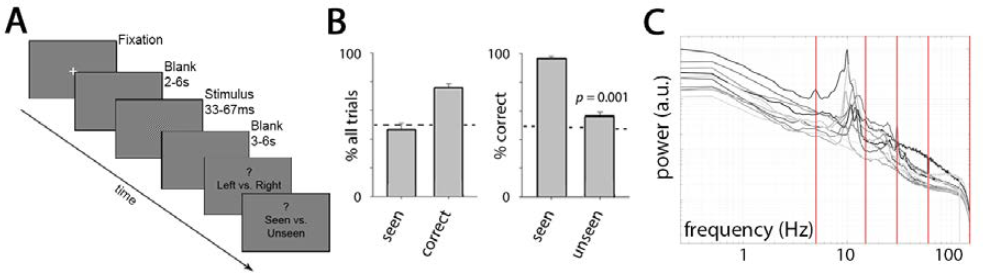
Task paradigm, behavioral results and MEG activity power spectrum. (**A**) In every trial, subjects discriminated the orientation of a brief (33-67 ms), low-contrast Gabor patch pointing randomly to upper left or upper right with equal chance, and then reported whether they consciously saw it or not. (**B**) *Left*: Percentage of trials in which subjects reported seeing the stimulus (‘seen’), and in which their orientation discrimination was correct (‘correct’). *Right*: %correct (determined by orientation discrimination) in seen and unseen trials separately. Results show mean and s.e.m. across 11 subjects. (**C**) Broadband MEG power spectrum averaged across all sensors for each of the 11 subjects (gray to black traces). Red lines correspond to 5, 15, 30, 60 and 150 Hz – boundaries for frequency-band-specific analyses.

### Transient, Fast-Evolving Neural Dynamics during Conscious Perception

Based on our previous findings from massive-univariate analyses [7], we first focused on the SCPs by extracting MEG activity in the 0.05 – 5 Hz range, which constitutes the low-frequency component of broadband, arrhythmic activity (Fig. 1C). To examine the temporal evolution of large-scale neural dynamics underlying conscious vs. unconscious processing, we investigated neural activity trajectory in its multi-dimensional state space [21, 23, 24, 31]. At a given time point, the state of neural activity (measured by whole-head MEG) can be described as a point in a high-dimensional space, where each axis is defined by the activity of one of the 273 MEG sensors. The concatenation of these points over time forms a “population activity trajectory”. Because of correlation between activity from different sensors, the dimensionality of the state space could be substantially reduced to a much lower dimension, where each axis is defined by a linear combination of sensors. Following earlier studies [21, 23, 25, 32], we applied PCA and defined the state space using the first few principal components (PCs). The top five PCs explained 72.2 ± 1.1% (mean ± s.e.m. across subjects) of total variance in the data (Fig. 2A; for the scalp distribution of PC coefficients, see Fig. 3A). Hence, all PCA-based results were computed using the top five PCs unless otherwise noted. As shown below, the reported results are robust against the choice of number of PCs, ranging from as few as 3 up to all 273.

**Fig. 2.**
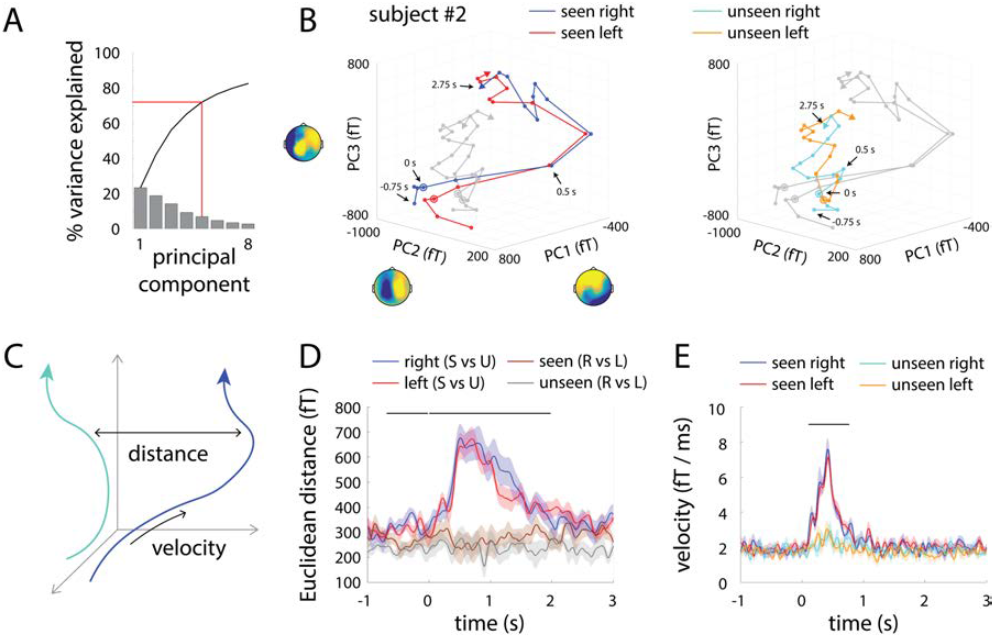
SCP (0.05 – 5 Hz) activity trajectories in seen and unseen trials. (A) Variance explained by each PC. The top 5 PCs explained > 70% variance and are used in subsequent analyses, unless otherwise noted. (**B**) Trial-averaged activity trajectories in the 3-dimensional PC-space from an example subject, for seen and unseen trials separately, under the presentation of the right-tilt (blue and cyan) or left-tilt stimulus (red and orange). Both plots show the same trajectories (i.e. gray trajectories in each plot are the same as colored trajectories in the other plot). For visualization, a 500-ms-length, half-overlapping moving average window was applied. Dots indicate the locations of each time point from 750 ms before to 2.75 sec after stimulus onset, with 250-ms steps. Circles indicate the time of stimulus onset. Topographical distribution of the PC coefficients is displayed adjacent to each axis. fT: femtotesla. (**C**) Schematic of Euclidean distance and velocity measurements. (**D**) Group-average pair-wise Euclidean distance between trajectories shown in B. Blue and red lines show the distance between seen and unseen trajectories under the presentation of the right-tilt (blue) or the left-tilt stimulus (red). Brown and gray lines show the distance between trajectories corresponding to different stimulus orientations under seen and unseen conditions, respectively. Horizontal bar indicates time points at which the distances between seen and unseen trajectories significantly exceed those between left- and right- tilt stimuli (2-way ANOVA, *p* < 0.05, cluster-based permutation test). (**E**) Group-average velocity of activity trajectories at each time point for different trial types shown in B. Horizontal bar indicates time points where seen and unseen trajectory velocities significantly differ (2-way ANOVA, *p* < 0.05, cluster-based permutation test). Shaded areas represent s.e.m. corresponding to the within-subject ANOVA design [65].

**Fig. 3.**
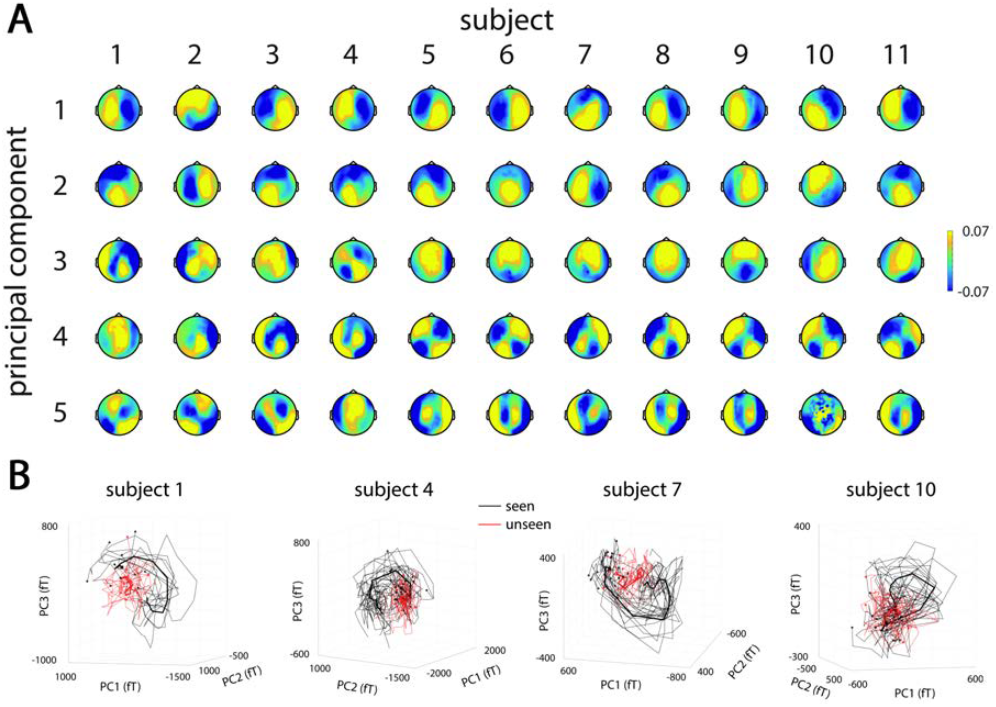
PC topographies and single-trial trajectories in the SCP range. (**A**) Topographical plots of the PC coefficients for the first 5 PCs in all 11 subjects. (**B**) Single-trial activity trajectories from four representative subjects. Each thin trace represents trajectory averaged across five randomly selected, unique trials within a condition (seen or unseen). Thick lines indicate the average across all seen (black) or unseen (red) trials. Trajectories are plotted from 1 sec before to 3 sec after stimulus onset; black and red dots indicate their origins. Seen and unseen trials follow qualitatively different trajectories at the single-trial level.

To obtain intuitions, we first visualized population activity trajectory by plotting the activity of the top three PCs, which together accounted for 56.2 ± 1.4% variance. Fig. 2B shows trial-averaged activity trajectories from a representative subject in seen and unseen conditions for the two stimulus orientations, respectively. A location in this three-dimensional space corresponds to the extent to which the spatial pattern encoded in each of the top three PCs (shown by the PC coefficient topography next to each axis) is present, and trajectories represent changes in the relative prominence of these patterns over time. Some qualitative observations can be made: First, seen and unseen trials follow distinct trajectories that start at different locations, while the trajectories corresponding to different stimulus orientations are very similar. Second, following stimulus input, the activity trajectories in seen trials move quickly to a distant location in the state space, whereas unseen trajectories remain closer to their starting point.

We used two quantitative measures to assess these observations. Importantly, data were analyzed in each subject’s individual PC space, and statistics (such as Euclidean distance and velocity, Fig. 2C) were pooled across subjects. *First*, we measured the Euclidean distance between trajectories in the state space (Fig. 2D). The distance between seen and unseen trajectories under the same physical stimulus (blue and red lines) was significantly greater than the distance between trajectories corresponding to different stimulus orientations but under the same subjective perceptual state (brown and gray lines) from 700 ms *before* stimulus onset to 2 sec after (Fig. 2D, black bar). In other words, as expected, the state of subjective awareness (seen vs. unseen) had a much greater effect on large-scale neural activity than the fine-grained feature of the stimulus (Gabor patch orientation). The time course of Euclidean distance between seen and unseen trajectories peaked around 500 – 750 ms following stimulus onset.

*Second*, we calculated the velocity of population activity trajectory for different conditions (Fig. 2E). In seen trials, activity trajectories accelerated drastically following stimulus onset, reaching peak velocity around 400 ms. Velocity returned to baseline at ~1 sec, far outlasting the duration of the visual stimulus (33 – 67 ms). By contrast, unseen trials exhibited a much smaller increase in velocity after stimulus onset, and the difference in velocity between seen and unseen trajectories is highly significant (Fig. 2E). These results confirm the above impression that large-scale SCP activity evolved more quickly in seen trials following stimulus input. Moreover, the separation between seen and unseen trajectories reached peak at ~500 ms (Fig. 2D), when trajectory velocity is near its peak (Fig. 2E); this suggests that neural activity distinguishing seen from unseen trials is fast changing (i.e., transient) instead of being in a steady state. We note that this analytical approach for adjudicating between steady-state vs. transient activity was previously applied in an animal model [23].

### Control Analyses for State-Space Findings

Since the above analyses were calculated on trial-averaged trajectories in each condition, could it be that unseen trials were more variable, and trial-averaging artificially reduced the velocity of unseen trajectories? To test this possibility, we computed velocity using single-trial trajectories (Fig. 4B). This analysis confirmed the original finding (Fig. 2E, reproduced in Fig. 4A): Following stimulus input, velocity of population activity trajectory increased significantly more in seen trials than unseen trials. In addition, all of the above results remained similar when the top three (explaining 56.2 ± 1.4% variance) or top eight (explaining 82.7 ± 0.8% variance) PCs were used in the analyses, as well as when the top 21 PCs (explaining 95.2 ± 0.4% variance) or all 273 PCs were used (Fig. 5).

**Fig. 4.**
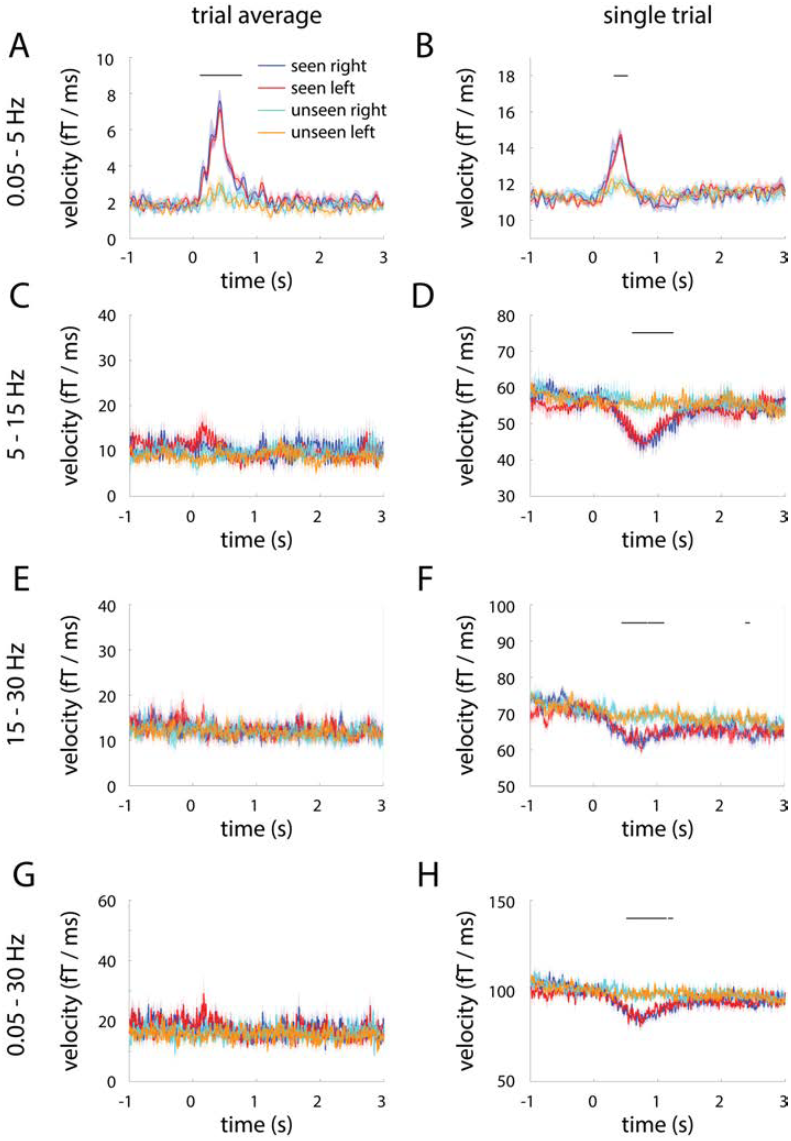
Trial-averaged and single-trial trajectory velocities in different frequency bands. (A, C, E, G) Trial-averaged trajectory velocity in the 0.05 – 5 Hz (A), 5 – 15 Hz (C), 15 – 30 Hz (E) and 0.05 – 30 Hz (G) band in seen and unseen trials, split according to stimulus orientation. **(B, D, F, H)** Single-trial trajectory velocity in the same frequency ranges. Horizontal bars indicate time points where seen and unseen velocities significantly differ (2-way ANOVA, *p* < 0.05, cluster-based permutation test). Shaded areas represent s.e.m. corresponding to the within-subject ANOVA design.

**Fig. 5.**
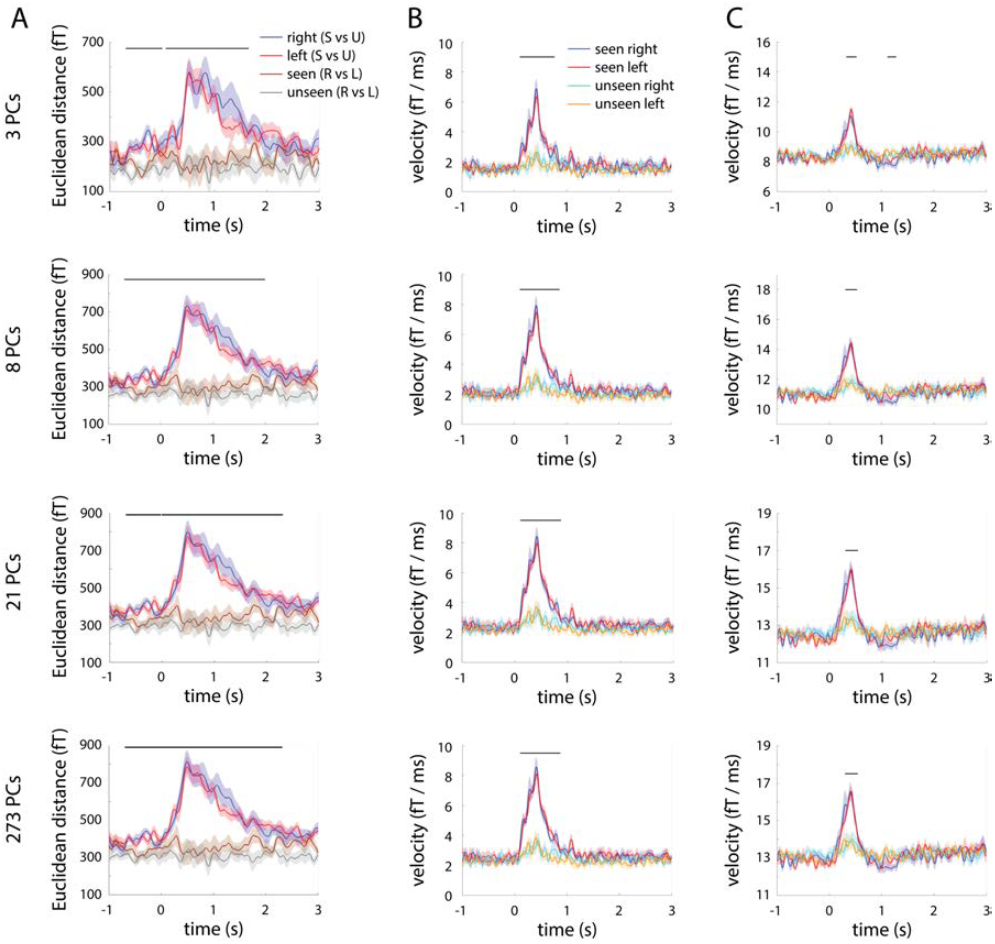
Control analyses showing the robustness of SCP trajectory results to the number of PCs included in the analysis. (**A**) Same as Fig. 2D, but using 3, 8, 21, or 273 PCs. (**B-C**) Same as Fig. 4A-B, but using 3, 8, 21, or 273 PCs.

Given that PCA was computed across all trials, could seen trials contribute more to the top few PCs, and hence have an unfair advantage in the comparison? To address this question, we conducted a control analysis where PCA was performed using only seen trials, or only unseen trials, and the extracted PC coefficients were then applied to all trials. This analysis showed that using the PCs defined by seen trials alone (Fig. 6D-F, top) or unseen trials alone (Fig. 6D-F, bottom) reproduced the above results. Indeed, the correlation structure amongst sensors (and hence, the PC decomposition) was very similar between seen and unseen conditions (Fig. 6A-C), which likely reflects the fact that the amplitude of the ERFs evoked by seen and unseen stimuli (~10^−14^ – 10^−13^ T) are much smaller than that of ongoing MEG activity (~10^−12^ T or higher). In other words, the PC decomposition captures intrinsic correlation structure amongst sensors that is minimally altered by task condition.

**Fig. 6.**
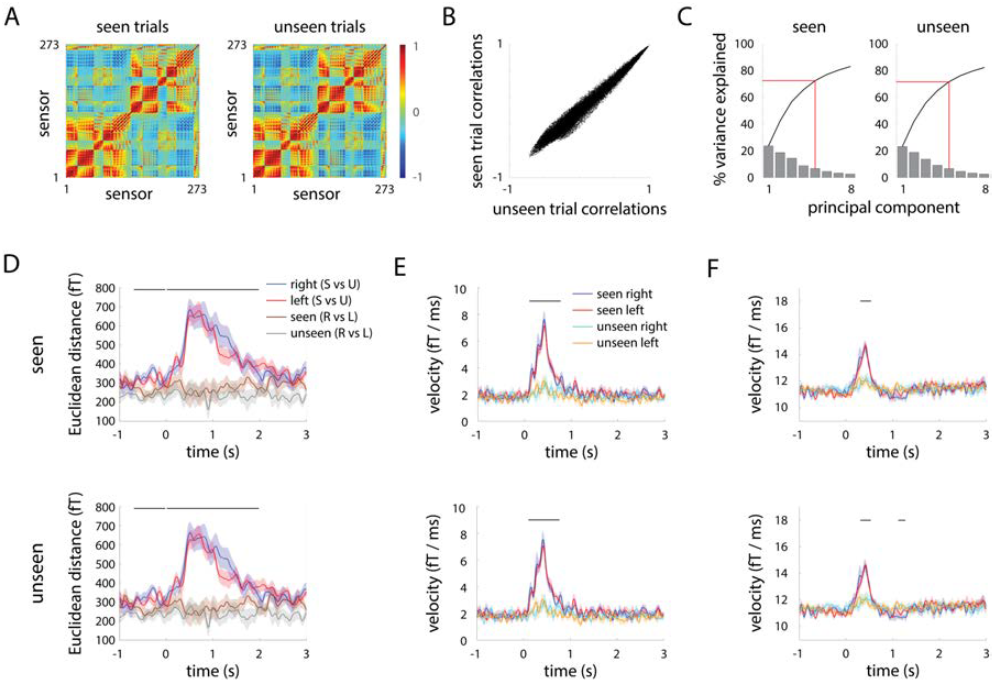
Control analysis showing that SCP trajectory results are robust to PCA decomposition method. (**A**) Sensor-by-sensor correlation matrix computed using temporally concatenated seen (left) or unseen (right) trials from an example subject. These correlation matrices were used for PCA decomposition. (**B**) Element-by-element scatter plot of the correlation matrices in A. (**C**) Variance explained by each PC when PCA was conducted on seen trials (left) or unseen trials (right) alone. The top five PCs explain > 70% of variance in both cases and are used for subsequent analyses. (**D-F**) Euclidean distance (D), trial-averaged trajectory velocity (E), and single-trial trajectory velocity (E). Top: PC coefficients (i.e., loading matrix) extracted from seen trials alone are applied to both seen and unseen trials. Bottom: PC coefficients extracted from unseen trials alone are applied to both seen and unseen trials. All analyses use 0.05 – 5 Hz activity.

A final control analysis confirmed that the horizontal and vertical eye movement components extracted from the ICA (and removed from data before analyses) did not differ between the seen and unseen conditions, nor did the saccade rate estimated from these components, suggesting that the results could not be attributed to any potential difference in eye movements between perceptual conditions.

### Stabilization of High-Frequency Activity during Conscious Perception

We next probed the specificity of our findings by investigating both trial-averaged and single-trial trajectory velocities in higher frequency bands. For activity filtered in the 5 – 15 Hz range (extracting the alpha oscillation; see Fig. 1C), the velocity of trial-averaged trajectories fluctuated around baseline following stimulus onset, and there was no significant difference between seen and unseen conditions (Fig. 4C). Interestingly, single-trial trajectories in the 5 – 15 Hz range exhibited a *decrease* in velocity from ~250 ms to 1.25 sec following stimulus onset in seen trials only (Fig. 4D), suggesting that population activity in this frequency range becomes more stable when the stimulus is consciously perceived. This pattern of results in seen trials is consistent with reduction of power and a lack of phase-locking across trials at individual-sensor level in this frequency range [7]. The 15 – 30 Hz band (extracting the beta oscillation; Fig. 1C) exhibited qualitatively similar results (Fig. 4E-F). There was no significant difference in velocity between seen and unseen conditions in higher-frequency ranges (30 – 60 and 60 – 150 Hz; low- and high-gamma activity, respectively) for either trial-averaged or single-trial trajectories. Together, these results suggest that the transient dynamics in seen trials is specific to the SCP range.

To examine the sum total effect of different frequency ranges exhibiting a difference between seen and unseen trajectory velocities (0.05 – 5 Hz, 5 – 15 Hz, 15 – 30 Hz), we investigated trial-averaged and single-trial trajectory velocities for the 0.05 – 30 Hz band. The results are similar to that of 5 – 15 Hz and 15 – 30 Hz activity (Fig. 4G-H). This suggests that trajectory velocity in this relatively broad frequency band is dominated by contributions from its high-frequency range, presumably because baseline velocity is much larger for higher frequency bands.

### Seen and Unseen Trajectories Are Well Separated both *Before* and After Stimulus Input

To explore the distribution of single-trial data, first, pseudo-single-trial trajectories (averaged over 5 randomly selected single trials) in seen and unseen conditions are plotted for four representative subjects in Fig. 3B, and it can be observed that they occupy separate regions of the state space throughout the trial. Next, we quantitatively assess the separation between seen and unseen activity trajectories at the single-trial level by performing single-trial decoding analysis. Strikingly, using SCP activity from all sensors, the classifier predicted subjective perceptual outcome (“seen” vs. “unseen”) significantly above chance at every time point from 1 s *before* to 3 s after stimulus onset (*p* < 0.05, cluster-based permutation test, Fig. 7A, magenta; a classifier constructed using 273 PCs had nearly identical performance, see Fig. 8A). The classifier reached peak performance at 700 ms following stimulus onset with an accuracy rate of 80 ± 2%. Importantly, the classifier’s performance was well above chance in the 1 s period *before* stimulus onset. The amplitude envelope of the SCP yielded weaker but significant decoding of subjective perception in the post-stimulus period, while its pre-stimulus decoding performance fell to near-chance level (Fig. 7A, green). Hence, SCPs’ contribution to decoding subjective perception in the pre-stimulus period depends not on its power, but on the moment-to-moment fluctuations of its activity.

**Fig. 7.**
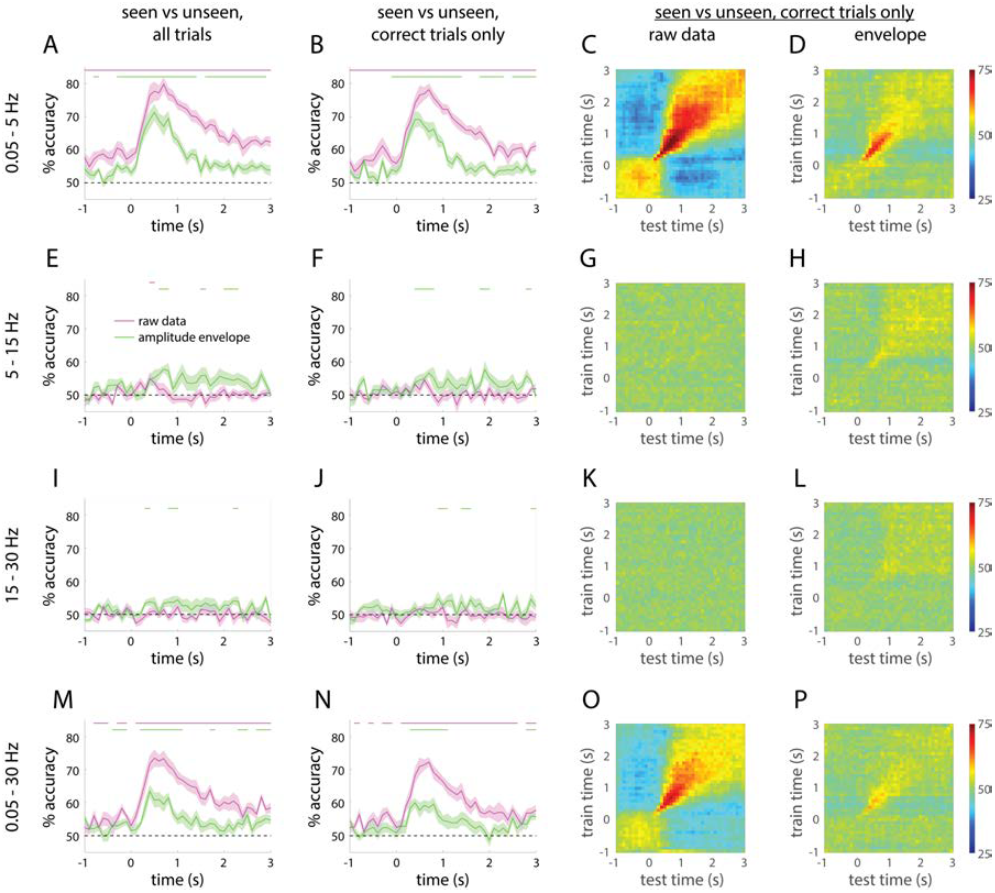
Single-trial decoding of seen vs. unseen perceptual outcome. (**A**) Single-trial classification accuracy using SCP activity across 273 sensors of seen vs. unseen perceptual outcome (magenta). Decoding accuracy is significantly above chance at every time point throughout the epoch (magenta bar). Classification accuracy is attenuated when using the amplitude envelope in this frequency range (green). **(B)** Similar classification accuracy is found when restricting the analysis to correct trials. **(C-D)** Temporal generalization of decoding results in A for filtered SCP activity (C) and its amplitude envelope (D), respectively. Rows indicate time points used to train the classifier and columns indicate time points used for testing. **(E-H)** Same as A-D, but for 5 – 15 Hz data. **(I-L)** 15 – 30 Hz data. **(M-P)** 0.05 – 30 Hz data. Horizontal bars indicate time points where decoding accuracy is significantly above chance (*p* < 0.05, cluster-based permutation test). Shaded areas represent s.e.m. across subjects.

**Fig. 8.**
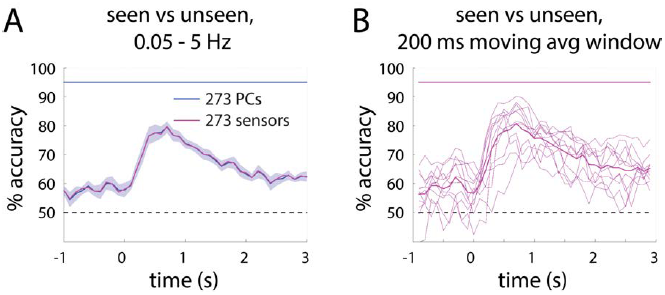
Seen vs. unseen decoding results using PCs and moving-average windows. (**A**)The magenta trace is reproduced from Fig. 7A, showing the accuracy of single-trial classification of seen vs. unseen perceptual outcome, using filtered SCP activity across 273 sensors. Blue trace plots seen vs. unseen decoding accuracy when the classifier was constructed using activity across 273 PCs; shaded area shows s.e.m. across subjects. The accuracy obtained using 273 PCs is significantly above chance at every time point from 1 sec before to 3 sec after stimulus onset (blue horizontal bar, *p* < 0.05, cluster-based permutation test). (**B**) Seen vs. unseen decoding result using a moving-averaging window (200-ms-length, half-overlapping) applied to the full-band data. Thin traces are the decoding results from each of the 11 subjects; thick trace is the group average. Magenta horizontal bar shows that the result is significant at every time point (*p* < 0.05, cluster-based permutation test).

One concern in interpreting the above results is that seen and unseen trials differ in aspects other than subjective perception of the stimulus, such as performance in the orientation discrimination task. To control for objective performance, we re-conducted the analysis using correct trials only, which yielded nearly identical results (Fig. 7B), suggesting that the decoding results were not driven by the categorical distinction between correct and incorrect trials (but see [33]). In addition, to ensure that the significant decoding result in the pre-stimulus period was not confounded by the (< 5 Hz) low-pass filter’s spreading of post-stimulus activity into the pre-stimulus period, we repeated the above analysis using full-band data. With a 200-ms-length, 100-ms-step sliding window, significant decoding result was obtained in every time point from 900 ms before to 2.9 sec after stimulus onset, with nearly identical accuracy to that achieved using filtered SCP activity (Fig. 8B, thick magenta trace), which confirms that pre-stimulus decoding was not affected by post-stimulus activity. Fig. 8B also shows robust decoding at the single-subject level (results from all 11 subjects are plotted in thin magenta traces).

To shed further light on the dynamical nature of neural activity underlying conscious perception, we conducted cross-time decoding analysis, which examines how the classifier trained at one time point generalizes to other time points [24, 34]. Using filtered SCP activity, classifier trained at a given pre-stimulus time point generalizes equally well to other pre-stimulus time points, as indicated by the uniformly colored square at the bottom left of Fig. 7C. This suggests that the representation format of predictive decoding of subjective perception is roughly constant in the pre-stimulus period. Starting from 200 ms following stimulus input, the temporal generalization of decoding narrows around the diagonal for about 1 sec, indicating rapidly changing serial computations – consistent with the high trajectory velocity in this time period shown in Fig. 4A-B. This was followed by a period of broad temporal generalization until the end of the trial, suggesting stable patterns of neural activity, which may reflect retention of stimulus judgment in working memory before response prompts. The cross-time decoding matrix using the amplitude envelope of SCP showed a diagonal pattern in the 200 ms – 1 sec period following stimulus onset, suggesting transient, serial computations triggered by stimulus input, followed by very weak and broad temporal generalization (Fig. 7D). Together, the above findings suggest that large-scale SCP activity is well separable between seen and unseen perceptual outcomes at the single-trial level, both before and after stimulus onset. In other words, depending on the initial state of large-scale SCP activity, sensory input from an identical physical stimulus sends the brain onto distinct trajectories of activity pattern evolution that correspond to consciously perceiving the stimulus or not.

By contrast, MEG activity filtered in the 5 – 15 Hz range did not yield above-chance decoding of subjective perception (Fig. 7E-F, magenta, and Fig. 7G). The amplitude envelope in the 5 – 15 Hz range provided marginally above-chance decoding of subjective perception (Fig. 7E-F, green, and Fig. 7H), likely due to stronger power reduction in this frequency range in seen trials [7]. Results from 15 – 30 Hz band are similar (Fig. 7I-L). Using the full band data ranging from 0.05 to 30 Hz, decoding performance was similar but inferior to that using the 0.05 – 5 Hz activity (Fig. 7M-P). This indicates that MEG activity in the 5 – 30 Hz range effectively functioned as a source of noise rather than signal for the decoding of subjective perception. We found a failure of above-chance decoding for seen vs. unseen trials in the 30 – 60 and 60 – 150 Hz range using either the filtered activity or its amplitude envelope.

We further examined decoding of other task parameters using SCPs across all sensors, which revealed i) non-significant decoding of objective performance (correct vs. incorrect) when controlling for subjective awareness by analyzing unseen trials only (Fig. 9A-B), and ii) transient above-chance decoding of stimulus orientation and discrimination response (left vs. right) in seen trials (Fig. 9C-D), but not unseen trials (Fig. 9E-F).

**Fig. 9.**
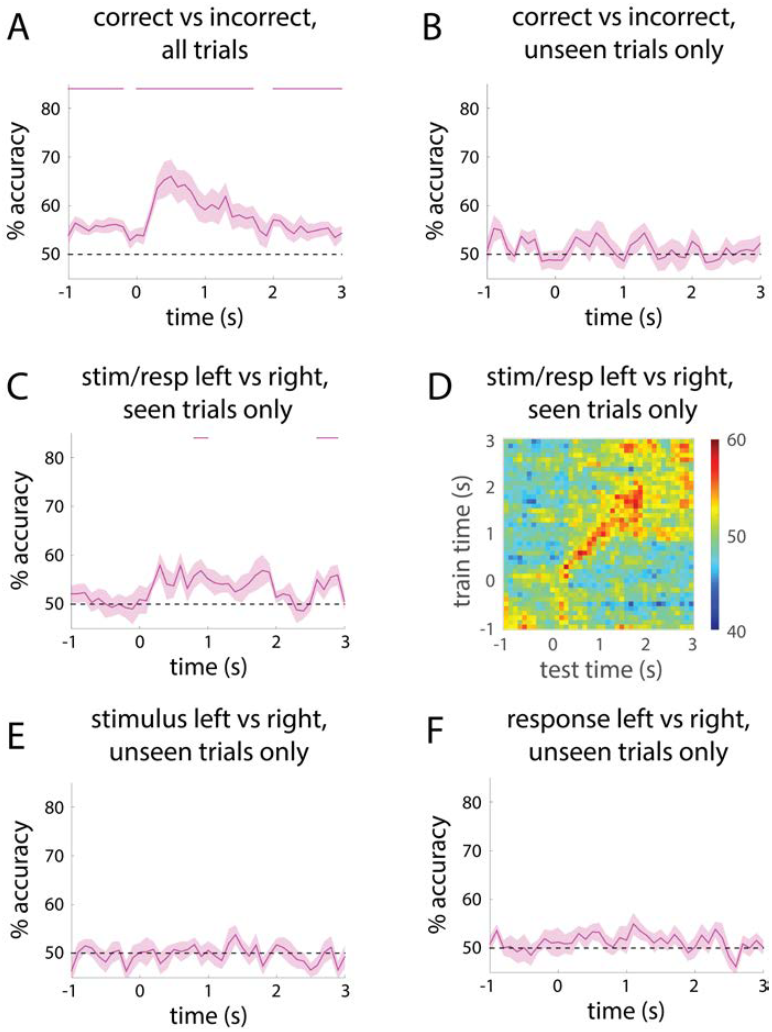
Additional decoding results using SCP activity across all sensors. (**A**) Decoding of correct vs. incorrect orientation discrimination performance. (**B**) Decoding of correct vs. incorrect performance for unseen trials only. (**C**) Decoding of stimulus orientation / discrimination response (left vs. right) for seen trials only. Note that due to the removal of a very small number of seen & incorrect trials, seen trials were always correct; hence, stimulus orientation and discrimination response are identical for analyzed seen trials. (**D**) Temporal generalization of decoding result in C. Rows indicate time points used to train the SVM classifier and columns indicate time points used for testing. (**E-F**) Decoding of stimulus orientation (E) and discrimination response (F) for unseen trials only. Shaded areas represent s.e.m. across subjects and horizontal bars show time points where decoding accuracy is significantly above chance (*p* < 0.05, cluster-based permutation test).

What is the spatial distribution of MEG activity that discriminates between seen and unseen trials? A simple way to visualize this is to plot the grand-average MEG activity topography separately for seen and unseen trials (Fig. 10A). A more sophisticated – and directly relevant to our decoder – approach is to visualize the MEG activation pattern utilized by the SVM decoder [35]. The activation pattern is a transformation of the decoder weights (Fig. 10B), allowing for inferences regarding the neurophysiological sources contributing to the classification between seen and unseen trials (for details see Methods). The activation pattern maps averaged across subjects are shown in Fig. 10C, which suggest contribution from widespread cortices, especially frontal regions, to the pre-stimulus decoding result; and following stimulus input, a sequence of activity patterns starting from the occipital regions and extending into temporal and frontoparietal cortices.

**Fig. 10.**
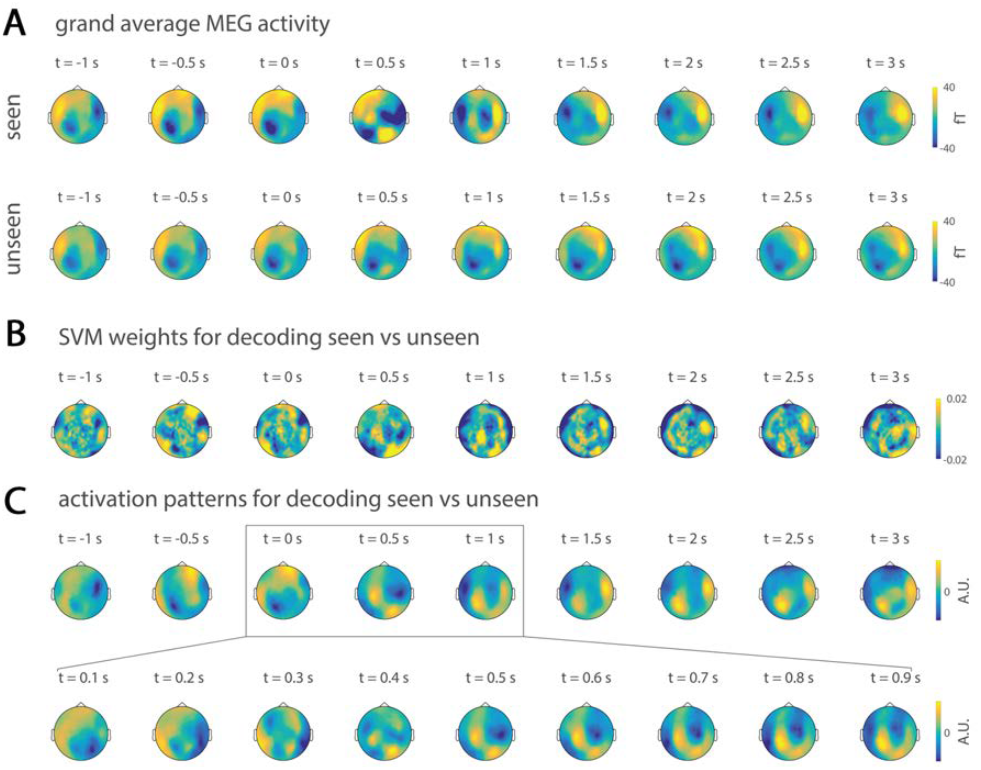
Topographical patterns of SCP activity distinguishing seen from unseen trials. (**A**) Grand-average SCP activity topography from 1 sec before to 3 sec after stimulus onset in seen and unseen conditions. (**B**) Topographic distribution of SVM decoder weights for the classification of seen vs. unseen perceptual outcome, averaged across subjects. (**C**) Activation patterns contributing to the seen vs. unseen classifiers, averaged across subjects.

### Control for Slow Attentional Fluctuations

The significant decoding of subjective awareness in the pre-stimulus period using SCPs (Fig. 7A-has two alternative, though not mutually exclusive, interpretations. It could reflect either fluctuations in the pre-stimulus attentional state [36] or spontaneous brain activity unrelated to attention [37]. Because in our task the stimulus was always presented at fovea and its occurrence time was largely unpredictable to the subject, there was no explicit modulation of spatial or temporal attention. An endogenous source of attentional fluctuation over the course of the experiment may be fluctuations in subjects’ tonic alertness or vigilance [38]. Given that tonic alertness fluctuates at a relatively slow time scale [39], we designed the following analysis to assess its contribution to our results. For each subject, we partitioned all trials into 3 groups using two methods: 1) an interleaving method in which every 3^rd^ trial was selected, which resulted in 3 groups of trials, each evenly distributed across the entire experiment; 2) a chunking method in which the first 1/3 of trials were selected into one group, the next 1/3 a different group, and the last 1/3 a third group (%seen rate was not significantly different between the three groups; F_2,20_ = 0.097, *p* = 0.9). We then constructed a SVM to decode the state of subjective awareness using each group of trials separately. If the pre-stimulus decoding result was contributed by slow attentional fluctuation’s modulation of conscious access, the interleaving method would capture more attentional variation across the experiment within each group of trials and thus produce higher decoding accuracy. However, decoding accuracy was nearly identical at each pre-stimulus time point for the two methods. Paired t-tests across subjects revealed only one time point (at -600 ms) with a significant difference (*p* <.05, uncorrected), with the interleaving method yielding worse performance than the chunking method, contrary to what would be expected if slow vigilance fluctuation contributed to decoding at this time point. These results support the interpretation that the significant pre-stimulus decoding of subjective awareness was unlikely to be driven by slow fluctuations in vigilance over the course of the experiment. Nonetheless, we cannot at present rule out the possibility that it had contributions from attentional fluctuations occurring at faster, trial-to-trial timescales [40].

### Transient Neural Dynamics Are Robust to Small Variations in the Initial State

Thus far, we have shown that transient dynamics in the SCP range characterizes conscious processing of a threshold-level visual stimulus, and that these dynamics follow a trajectory that is distinct from that during unconscious processing. Moreover, which trajectory is followed by neural dynamics in a given trial depends to a large extent on the initial state of large-scale SCP activity. These results raise an important question: Are transient neural dynamics during conscious processing robust to noise? Robustness against noise is important for information transmission and computing, and theoretical studies have shown that recurrent neural networks can be trained to exhibit robust, transient dynamics that are resistant to noise [41–43]. On the other hand, since these transient dynamics manifest drastic acceleration (Fig. 4A-B), one might expect that as velocity increases, noise in the system could become amplified and render the dynamics unstable, as seen in recurrent neural networks exhibiting chaotic dynamics [41, 44]. We investigated the robustness of population activity trajectories during conscious or unconscious processing of the visual stimulus using measures of across-trial variability. If the transient neural dynamics are unstable, across-trial variability should increase with time. Conversely, variability reduction would suggest that the dynamics are resistant to small variations in the initial state, hence robust.

We first computed across-trial variability of SCP for each MEG sensor. In seen trials, stimulus onset induced a reduction in across-trial variability in many sensors, whereas the extent of variability reduction was weaker for unseen trials (Fig. 11A-C). To describe across-trial variability at the large-scale population activity level, we estimated the volume of state space occupied by the distribution of single-trial trajectories by calculating the across-trial variability (s.d.) of each PC and multiplying it across the top five PCs (see Methods). Because PCs are orthogonal, this product reflects the volume of activity distribution in the state space. The volume decreased massively following stimulus onset in seen, correct trials, reaching 47.7 ± 6.0% reduction at 715 ms (Fig. 11D). Volume reduction was smaller in unseen trials, reaching 18.0 ± 6.4% at 657 ms for unseen, correct trials and 27.1 ± 10.4% at 377 ms for unseen, incorrect trials. Importantly, even though the visual stimulus was very brief (lasting 33 – 67 ms), volume reduction in seen trials peaked much later (at 500 – 750 ms) and slowly returned to baseline over a 2.5-sec period, displaying a time course similar to that of the Euclidean distance between seen and unseen trajectories (Fig. 2D). Thus, a brief, threshold-level visual stimulus that is consciously perceived sends the brain onto a fast-changing trajectory that exhibits reduced across-trial variability over a ~2 sec period. This result suggests that neural activity during conscious processing belongs to the class of robust, transient dynamics that, while fast-changing, are resistant to small variations in the initial state [28, 41]. However, when the variation in the initial state is large enough, population activity will follow an entirely different trajectory that corresponds instead to unconscious processing (Figs. 2 & 7).

**Fig. 11.**
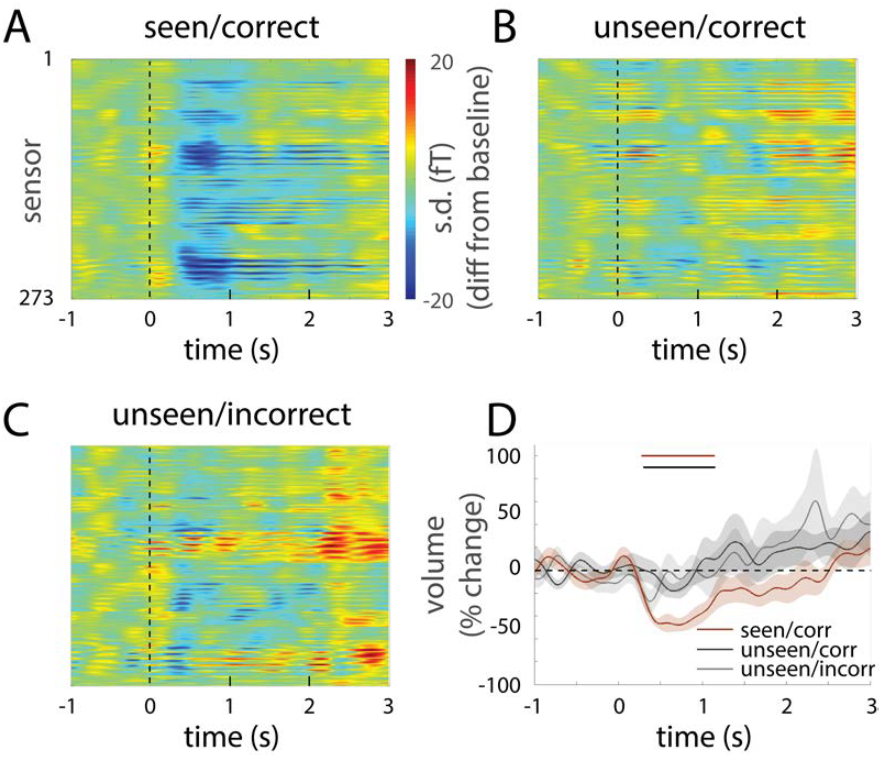
Across-trial variability reduction in the SCP range during seen trials. (**A-C**) Across-trial variability (s.d.) time course in all sensors for seen & correct (A), unseen & correct (B), and unseen & incorrect (C) conditions, measured as the difference between s.d. at each time point and the mean of the pre-stimulus period. (**D**) %change of state-space volume as compared to the pre-stimulus period. Brown horizontal bar indicates that in seen & correct condition, the change from baseline was significant from 322 ms to 1087 ms (one-sample t-test, *p* < 0.05, cluster-based permutation test). Black horizontal bar indicates that variability reduction was significantly greater for seen & correct than for unseen & correct condition from 385 ms to 955 ms (paired t-test, *p* < 0.05, cluster-based permutation test). Shaded areas represent s.e.m. across subjects.

### Norm vs. Angle of Population Activity Trajectory

The location of population activity in the state space at a given time point can be described by the combination of two variables – angle and norm of a vector pointing from the origin of the state space (where the value of each axis is 0) to this location (Fig. 12E, for details see Methods). Angle describes the direction in the state space that the vector points to and reflects the *relative* pattern of activity across axes (defined as sensors or PCs). Norm, the length of this vector (calculated as root mean square across the axes), reflects the total energy of activity. To shed further light onto the above findings, we next investigated the respective contribution of population activity norm vs. angle to decoding and across-trial variability results in the SCP range. Specifically, we asked: 1) Given that seen and unseen trajectories occupy separate regions of the state space throughout the trial (Fig. 7A-B), are they distinguished by their total amount of energy (norm) or spatial pattern of activity (angle), or both? 2) Given that the volume of the state space occupied by single trials shrinks during conscious perception (Fig. 11D), is it because the total amount of energy (norm) or spatial pattern of activity (angle) becomes more consistent across trials?

**Fig. 12.**
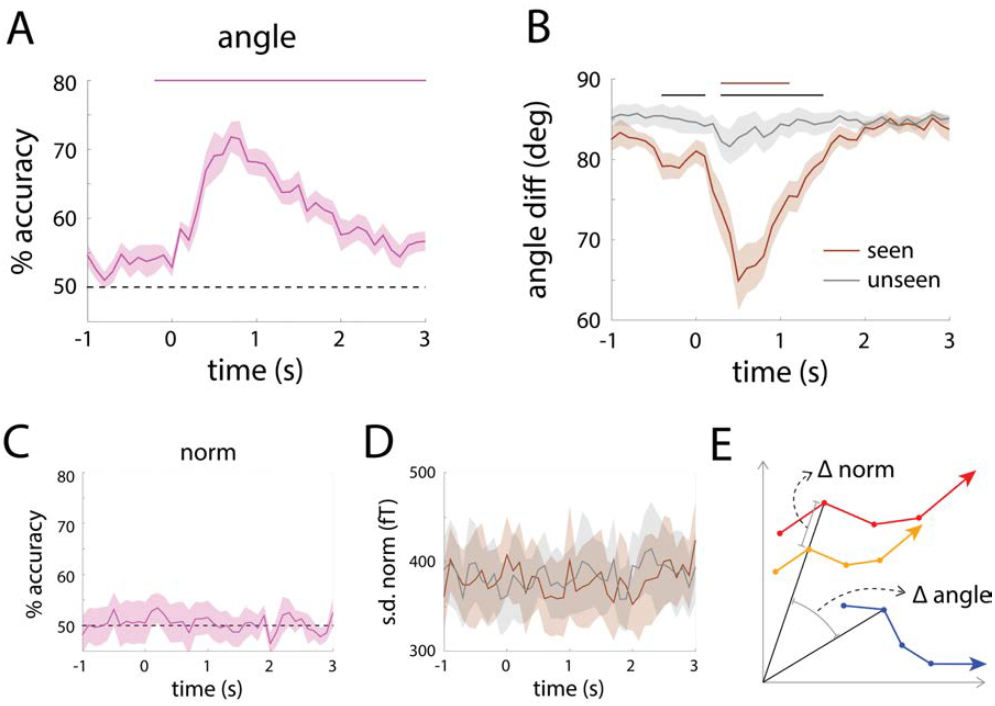
Single-trial decoding and across-trial variability using the angle (A-B) or norm (C-of population SCP activity. (**A**) Single-trial classification of seen vs. unseen perceptual outcome based on the angle was significantly above chance from 200 ms before to 3 s after stimulus onset (horizontal bar, *p* < 0.05, cluster-based permutation test). (**B**) Across-trial variability of angle in seen and unseen conditions. Brown bar indicates time points where angle variability for seen trials was significantly lower than the pre-stimulus baseline; black bar indicates time points where angle variability was significantly different between seen and unseen trials (*p* < 0.05, cluster-based permutation test). (**C**) Single-trial classification of seen vs. unseen perceptual outcome based on the norm was not significantly different from chance at any time point. (**D**) Across-trial variability (s.d.) of the norm in the seen or unseen condition did not exhibit any significant change relative to pre-stimulus baseline or difference between seen and unseen conditions. Shaded areas denote s.e.m. across subjects. (**E**) Schematic illustrating norm and angle in a 2-dimensional state space. The orange and red trajectories have the same angle at every time point (denoted by circles), but differ in norm. The orange and blue trajectories have the same norm at every time point, but differ in angle.

To address the first question, we performed single-trial decoding of subjective awareness using only the angle or the norm of population activity (defined by 273 sensors, for comparison with Fig. 7). Using angle alone, significant decoding was achieved at every time points from 200 ms before to 3 sec after stimulus onset (*p* < 0.05, cluster-based permutation test, Fig. 12A). By contrast, using norm alone, decoding accuracy fluctuated around chance (Fig. 12C). Hence, the fact that seen and unseen trajectories are well separable is due to their residing in different directions in the state space, while their distances to the origin are not significantly different. In other words, they are distinguished by their *relative* spatial pattern of activity across sensors, but not the total energy of activity.

To address the second question, we computed the across-trial variability of angle and norm, respectively. While across-trial variability of norm did not change significantly from the pre-stimulus baseline (Fig. 12D), across-trial variability of angle reduced substantially in seen trials following stimulus input, reaching significance from 300 to 1100 ms (*p* < 0.05, cluster-based permutation test, Fig. 12B, brown bar), and showed a small but non-significant decrease in unseen trials. Comparing seen and unseen conditions, across-trial angle variability was lower for seen trials in both the pre- and post- stimulus periods: from 400 ms before stimulus onset to 100 ms after, and then from 300 ms to 1.5 sec after stimulus onset (Fig. 12B, black bar). These results suggest that the stronger across-trial variability reduction accompanying conscious perception (Fig. 11D) was contributed by the angle of the population activity as opposed to the norm. In other words, the relative spatial pattern of activity, but not the total amount of energy, becomes more consistent across trials during conscious perception.

### Pre-stimulus Brain Activity Influences Unconscious Perceptual Decision Making

Lastly, we investigated whether the initial state of brain activity also influences subjects’ unconscious perceptual judgment about a fine-grained stimulus feature – the orientation of the Gabor patch – in unseen trials. Inspired by sampling-based Bayesian theory [45], we tested the following hypothesis: If pre-stimulus brain activity pattern is similar to that evoked by a given stimulus, the subject’s decision would be biased towards that same stimulus. To this end, we first computed the mean MEG activity for each sensor in a post-stimulus time window (varied from 100 to 500 ms length, time locked to stimulus onset) across trials for each subject, separately for the two stimulus orientations (Fig. 13A), which is referred to as the individual-specific “evoked templates”. Then, using the same window length (again locked to stimulus onset), we extracted the pre-stimulus MEG activity for each trial. Trials are sorted according to whether their pre-stimulus activity was more similar to the evoked template for left-tilt or right-tilt stimulus, using spatial correlation across sensors. For each group of trials, we employed signal detection theory [46] to calculate the bias of the subject’s responses (Fig. 13A, right). Given our interest in a fine-grained stimulus feature, we conducted this analysis using all MEG sensors, and separately using sensors covering each lobe in order to elucidate potential regionally-specific effects.

**Fig. 13.**
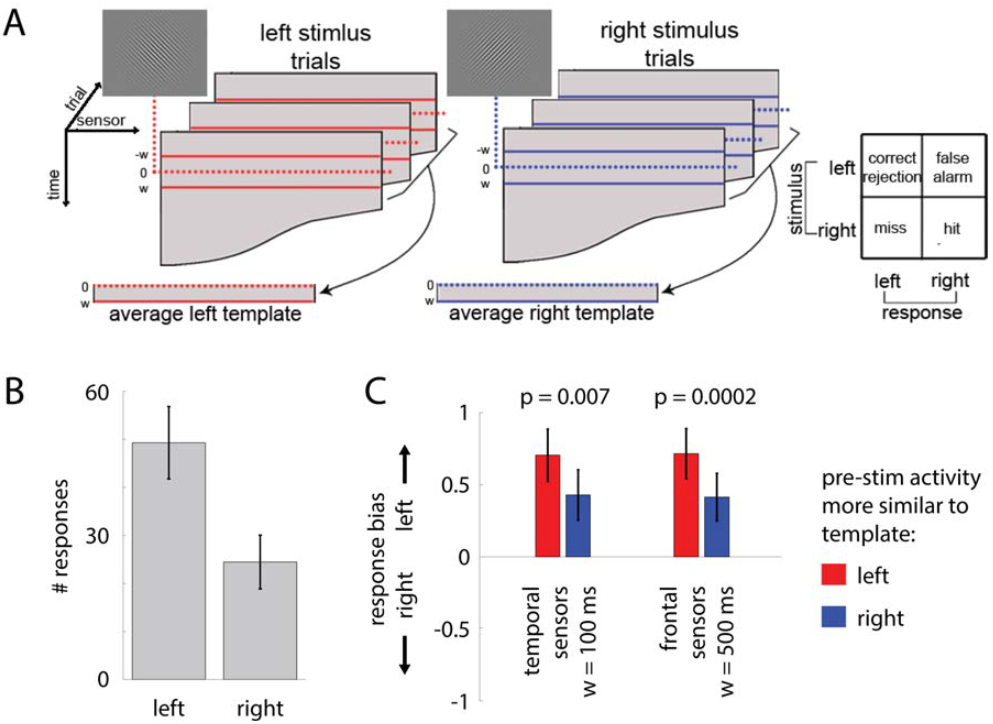
Pre-stimulus activity influences unconscious perceptual decision making. **(A**) Analysis stream. Stimulus-evoked activity templates for each subject were generated by averaging MEG activity in a post-stimulus time window of duration *w* across trials for each stimulus orientation. Mean pre-stimulus activity in a time window of the same duration was computed for each trial and compared to the two stimulus-evoked templates using spatial correlation. Trials were sorted into two groups by whether their pre-stimulus activity was more similar to the left-tilt or right-tilt stimulus template. A signal detection theory measure of response bias (c) was computed separately for each group of trials. The analysis was conducted using w = 100, 200, 300, 400, 500 ms, separately. (**B**) Across all trials, subjects tended to answer ‘left’ more often, suggesting that there was an innate response bias, presumably due to fixed stimulus-response mapping. (**C**) Results from temporal lobe sensors at *w* = 100 ms and from frontal lobe sensors with *w* = 500 ms. Error bars represent s.e.m. across subjects.

In this task, subjects had an innate response bias for answering the left stimulus (Fig. 13B), presumably due to fixed stimulus-response mapping (“left” response mapped to the more dominant index finger, while “right” response used the middle finger). Nonetheless, we found that the similarity between pre-stimulus activity and evoked template influenced response bias. With a window length of 100 ms, this effect was evident in temporal lobe sensors (*p* = 0.007), while a window length of 500 ms revealed an effect in frontal sensors (*p* = 0.0002, Fig. 13C). Both findings are significant after false-discovery rate correction across all tests conducted and are consistent with our hypothesis: If pre-stimulus activity is more similar to that evoked by the left stimulus, subjects were even more likely to answer “left”. The longer time window needed for the effect to manifest in frontal than temporal sensors accords with the idea of a gradient of temporal integration windows across the cortical hierarchy [47, 48]. These results demonstrate that pre-stimulus brain activity pattern influences subjects’ decision criterion in unconscious perceptual decision making about a fine-grained stimulus feature.

## DISCUSSION

In summary, we observed that depending on its initial state, large-scale slow cortical dynamics followed distinct trajectories that strongly predicted whether the subject consciously perceived a threshold-level visual stimulus or not (for a schematic summarizing our findings, see Fig. 14). The activity trajectories in the SCP range were well separated between seen and unseen trials from 1 sec before to 3 sec after stimulus onset with significant single-trial decoding throughout. By contrast, large-scale activity in higher-frequency ranges yielded chance-level or near-chance decoding performance. In addition, in seen trials, stimulus input triggered drastic acceleration of population activity trajectory in the SCP range, and the largest separation between seen and unseen trajectories occurred around 500 ms when trajectory velocity is highest, suggesting that neural activity underlying conscious perception is fast changing (“transient”) rather than steady. Furthermore, these transient SCP dynamics are robust to small variations in the initial state, as shown by the substantial reduction of across-trial variability following stimulus input. Yet, if the variation in the initial state is large enough, an entirely different trajectory would be followed that corresponds instead to unconscious processing. Together, these results shed light on the brain mechanisms governing conscious access and the nature of neural activity underlying conscious processing, which can be parsimoniously explained within an integrated framework of initial-state-dependent, robust, transient neural dynamics embodied in large-scale slow cortical potentials.

**Fig. 14.**
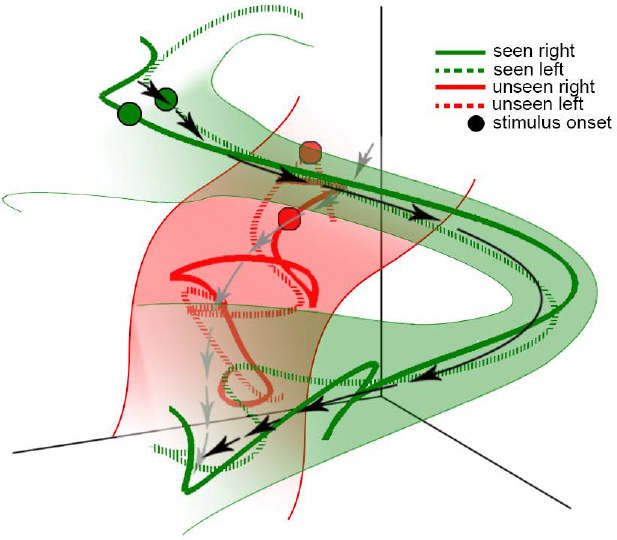
A schematic summarizing our results. Trajectories of seen trials (examples shown as green lines) begin in a location distinct from those of unseen trials (examples shown as red lines). The time of stimulus onset is indicated by circles, and different stimulus orientations are indicated by solid or dashed lines. For seen trials, the voyage of the trajectory through state space is characterized by marked increase in velocity following stimulus onset, indicated by the length of black arrows. Unseen trials, on the other hand, only accelerate minimally following stimulus onset, indicated by the length of gray arrows. Seen and unseen trajectories remain well separated throughout the trial. For seen trials, across-trial variability decreases substantially following stimulus onset (shown as green shading), suggesting that the trajectories are robust to small variations in the initial location.

Our results fit well with a class of theoretical models that emphasize the computational power of transient neural dynamics (‘trajectories’), whereby computation arises from the voyage through state space rather than the arrival at a fixed location [26, 28, 49]. Transient neural dynamics have been observed in local neuronal circuits in animal models [22–24, 31, 50]. Here we report, for the first time, that transient neural dynamics at the large-scale network level, embodied in the evolving spatial patterns of slow cortical dynamics, differentiate conscious from unconscious processing of an identical stimulus in humans. Thus, transient dynamics may be a neural computational mechanism that transcends spatial scales and species, and future theories on conscious processing should accommodate such robust, transient neural activity. Lastly, our finding that large-scale SCP dynamics during conscious processing is both robust to small variations in the initial state and sensitive to large variations thereof is nontrivial. Such coexistence of robustness and sensitivity could be implemented by the simultaneous presence of two robust trajectories in the state space, each with its own ‘energy basin’ (Fig. 14).

We further observed that pre-stimulus SCP activity influences subjects’ decision criterion when discriminating stimulus orientation in unseen trials (Fig. 13). To our knowledge, this is the first demonstration that spontaneous brain activity influences unconscious perceptual decision about a fine-grained stimulus feature. Consistent with sampling-based Bayesian theory [45], we found that if the pattern of pre-stimulus activity is more similar to that evoked by a given stimulus, subjects would be more inclined to select that stimulus.

The present result showing that conscious perception of a brief, threshold-level visual stimulus induces widespread and long-lasting across-trial variability reduction adds to the growing literature on stimulus-induced variability reduction (e.g., Churchland et al., 2010; He, 2013; He and Zempel, 2013; Schurger et al., 2015). Partitioning across-trial variability by the contribution of norm vs. angle revealed that variability reduction was mainly contributed by population activity angle (Fig. 12B), in accordance with Schurger et al. (2015). Interestingly, Schurger et al. also reported higher stability of neural activity over time in seen trials relative to unseen trials, whereas a seemingly conflicting finding was reported by Salti et al. (2015), who showed that the decoding of stimulus location had weaker temporal generalization in seen than unseen condition, indicating that neural activity in seen trials was less stable over time. Our results offer a reconciliation of this apparent discrepancy in the literature. Both of these previous studies used relatively broad frequency ranges (from DC to over 100 Hz and 30 Hz, respectively), but assessed neural stability with different measures – Schurger et al. used directional variance, a measure of the within-trial stability over time similar to our measure of single-trial trajectory velocity, whereas Salti et al. used temporal generalization of decoding performance. We found that decoding based on full-band data is sensitive to the transient dynamics in the SCPs (Fig. 7), while single-trial trajectory velocity based on full-band data is sensitive to stabilization in the high-frequency range (Fig. 4). The differential sensitivity of these measures to the spectral content of the MEG signal could account for why these previous studies yielded different conclusions about neural stability during conscious perception. Importantly, the present data elucidate that it is the transient dynamics embodied in the SCPs, not high-frequency stabilization, that provides robust single-trial decoding between seen and unseen perceptual outcomes.

An enduring debate in the pursuit of the neural basis of conscious perception concerns its localization in space and time. In the temporal domain, multiple studies have shown that late-onset (> 200 ms) activity correlates strongly with conscious perception [7, 9, 12, 13], but the timing of this activity could be modulated by prior expectation [51] and very early (~100 ms) correlates of conscious perception have also been reported [52, 53]. Importantly, many of these previous studies investigated event-related fields/potentials (ERFs/ERPs) that had the pre-stimulus baseline removed. The present results, describing continuous activity trajectories spanning pre- and post-stimulus periods, raise the possibility that instead of a well-circumscribed temporal period, the distinguishing feature of the neural dynamics underlying conscious processing is the evolution of its trajectory through state space over time (or equivalently, the activity pattern evolution). Depending on its initial state, which may reflect cognitive state (such as attention or expectation) and spontaneous activity fluctuations [54], neural activity may reach a desired state – a particular region in the state space – at different post-stimulus latencies. Future studies investigating ERFs/ERPs without baseline correction (such as Fig. 10A herein) could provide further linkage between the present findings and previous results using massive univariate analyses.

In the spatial domain, both higher-order frontoparietal areas and ventral visual areas have been emphasized [15, 16, 55–57]. Our results provide a potentially unifying picture across these previous findings. Sensor-level topography shows that both pre-stimulus activity in frontal regions and post-stimulus activity in occipitotemporal regions are more pronounced in seen than unseen trials (Fig. 10). Together with the cross-time decoding result showing sustained activity pattern in the pre-stimulus period and serial processing in the ~1-sec post-stimulus window (Fig. 7C), these results may suggest that both a pre-stimulus state of perceptual readiness and post-stimulus activation of visual areas jointly determine whether a stimulus achieves conscious access, in accordance with a previous proposal [58]. Importantly, the present results reveal the dynamical and frequency characteristics of these activities.

Our study illustrates that given the high degree of connectedness across the cerebral cortex [59, 60], analysis of distributed spatial patterns of activity complements traditional region-by-region analysis in the investigation of the neural mechanisms of conscious perception. We note that such an approach has also been applied to brain-wide neuronal dynamics in the zebrafish to reveal behaviorally relevant activity trajectories [32]. Future studies testing the generalization of the present findings using other task paradigms (such as manipulating stimulus duration and subjects’ attentional state) will be informative. For example, it will be interesting to know whether stabilization of large-scale SCP activity might be observed for stimuli with longer presentation durations.

In conclusion, we found that robust, transient, large-scale cortical dynamics in the SCP range (< 5 Hz) encode conscious visual perception. These neural dynamics exhibit both robustness and sensitivity to the initial state, such that small variations in the initial state do not prevent the dynamics from converging onto a robust trajectory, but large variations may switch the dynamics onto an entirely different trajectory that corresponds instead to unconscious processing. These results shed new light on the brain mechanisms governing conscious access and the neural dynamics underlying conscious processing, and account for them in a unified framework. Lastly, pre-stimulus brain activity not only influences conscious access of a stimulus but also impacts subjects’ unconscious perceptual decision making.

## METHODS

### Subjects

Subjects (N=11, 6 females, mean age 27, age range 22 to 38) were right-handed, neurologically healthy, and had normal or corrected-to-normal vision. All subjects provided written informed consent. The experiment was approved by the Institutional Review Board of the National Institute of Neurological Disorders and Stroke (protocol #14-N-0002).

### Stimuli and Task

Stimuli were presented on a Panasonic DLP projector with a 60-Hz refresh rate, onto a screen 75 cm away from the subject’s eyes. An optical filter was placed on the lens to reduce luminance of the stimulus so that every subject could reach a subjective threshold of stimulus duration longer than 16.7 ms – the limitation of the projector refresh rate. All subjects were dark-adapted for at least 30 min before the experiment began.

Each trial started with a white fixation cross on a gray background (Fig. 1A). When the subject was ready, s/he pressed a button to start the trial. After a blank screen of a random duration between 2 and 6 sec (following an exponential distribution), a Gabor patch (1° visual angle/cycle) with a very low contrast (1%) was presented for a short duration (exact stimulus duration was set individually for each subject in order to control subjective perception rates; see below). The orientation of the Gabor patch was randomly selected between 45° and 135° with equal chance. Then another blank screen with a duration randomly chosen between 3 and 6 sec (following an exponential distribution) was presented. The luminance of the blank screens was equal to the background luminance of the stimulus screen. The first blank period ensured that the subject could not predict the onset of the stimulus. Each trial ended with three sequential questions: i) A forced alternative-choice – Was the Gabor patch pointing to upper-left (135°) or upper-right (45°)? ii) Did you see the stimulus (Gabor patch) or not? iii) On a scale of 1 to 4, how confident are you about your answer to question ii? Subjects indicated their answers to the questions via a fibreoptic key-pad (LumiTouch). In this study we are only interested in subjects’ answers to the first and second question, indicating their objective performance and subjective awareness, respectively. Stimulus orientation was well-matched for seen (50.9% ± 0.13% left-tilting) and unseen (50.3% ± 0.09% left-tilting) trials.

The experiment was conducted in two stages. In the first stage, the duration of the Gabor patch was adjusted using Levitt’s staircase method, until an individually titrated threshold for subjective awareness was found (i.e., the subject answers “seen” to question ii in about half of the trials). The distribution of threshold duration across subjects was as follows: six subjects, 33.3 ms; three subjects, 50 ms; two subjects, 66.7 ms. Once the threshold duration was determined, in the second stage of the experiment, trials were shown repeatedly with identical stimulus duration at the subject’s individual threshold while MEG signals were continuously recorded. Subjects performed these trials in sessions that were less than 12 min long and were allowed to rest between sessions. Because seen and unseen trials were compared within subject, the physical property of the stimulus was exactly matched between them (contrast: 1%; duration: individual threshold). The entire experiment (including dark adaptation and staircase) lasted under 3 hours for each subject. The present data were previously reported in Li et al. (2014); full details of the task paradigm are described therein.

### Data Acquisition

Experiments were conducted in a whole-head 275-channel CTF MEG scanner (VSM MedTech). Two dysfunctional sensors were removed from all analyses. MEG data were collected using a sampling rate of 600 Hz with an anti-aliasing filter at **<** 150 Hz. Before and after each recording session, the head position of the subject was measured using coils placed on the ear canals and the bridge of the nose. All MEG data samples were corrected with respect to the refresh delay of the projector (measured with a photodiode).

### Preprocessing

MEG data was preprocessed using FieldTrip (http://fieldtrip.fcdonders.nl) in MATLAB (MathWorks). Each recording session was demeaned, detrended, band-pass filtered between 0.05 and 150 Hz with a fourth-order Butterworth filter, and notch-filtered at 60 and 120 Hz to remove power-line noise. Independent component analysis (ICA) was performed on the continuous data to remove eye-movement, cardiac, and movement artifacts. Subsequently we applied a second set of fourth-order Butterworth filters in ranges of 0.05 – 5 Hz, 5 – 15 Hz, 15 – 30 Hz, 30 – 60 Hz, and 60 – 150 Hz to extract activity in these different frequency bands, and Hilbert transform was used to extract their amplitude envelope time courses. Finally, data were epoched from 1 sec before to 3 sec after stimulus onset, and remaining trials with artifacts were rejected manually. In SVM decoding analyses, data were downsampled to 10 Hz in order to save computation time. All statistical results using a time-point-by-time-point analysis were corrected for multiple comparisons using cluster-based permutation tests [61], as described below.

### Principal Component Analysis (PCA) and State-Space Analysis in the SCP Range

For each subject, MEG data filtered in the 0.05 – 5 Hz range were concatenated across all trials. Pearson correlation was calculated between all sensor pairs, resulting in a 273 x 273 correlation matrix. PCA was performed on this correlation matrix using the pcacov function in Matlab, and the resulting PC coefficients (i.e., the loading matrix) were used to construct activity of each PC in each trial [for a detailed explanation of PCA method, see Briggman et al. (2015)]. Using the covariance matrix instead of correlation matrix for PC decomposition yielded very similar results.

For each subject, PCs were averaged across trials of the same type, with trial types defined by the state of subjective awareness (seen vs. unseen) and the orientation of the stimulus. To visualize activity trajectories, the activity at each time point is plotted as a location in the 3-dimensional space defined by the top three PCs. At each time point, distance between trajectories of different trial types was calculated as the Euclidean distance in the five-dimensional space defined by the top five PCs. Velocity of a trajectory was calculated as the distance between adjacent time points (in the 5-D PC space) divided by the incremental time unit (Δt = 1.67 ms). Both trial-averaged trajectory velocities and single-trial trajectory velocities were calculated. We also carried out analyses using the top 3 or 8 PCs, as well as the top 21 PCs or all 273 PCs.

For velocity analyses, we computed a 2 (stimulus: left tilt / right tilt) x 2 (awareness: seen / unseen) repeated-measures ANOVA at each time point. Cluster-based permutation test was performed separately for the statistical effects of stimulus, awareness, and their interaction. Clusters were defined as contiguous time points where *p* < 0.05, and cluster summary statistic was computed as the sum of the F-values in each cluster. Null distributions of cluster statistics were derived by performing the ANOVA analysis on 1000 random permutations of the original data and extracting the maximal cluster statistic yielded by each permutation. Clusters in the original data were considered significant if their summary statistic exceeded the 95th percentile of the null distribution. The effect of stimulus or stimulus x awareness interaction did not yield any significant result after correction.

In a control analysis, PCA was performed on the 273 x 273 correlation matrix derived from concatenated seen trials or unseen trials alone, and the PC coefficients were applied to all trials. The reconstructed PC activity was subjected to the same state-space analyses as described above. This analysis confirmed that the original PCA decomposition was not driven unfairly by variance contributed by seen or unseen trials.

### Trajectory Velocity in Higher Frequency Bands

PCA decomposition was carried out for each of the other frequency bands (5 – 15 Hz, 15 – 30 Hz, 30 – 60 Hz, and 60 – 150 Hz) separately, using a similar method as that described above. For each frequency band, we chose the minimum number of PCs that explain > 70% of variance in the data (averaged across subjects). This yielded 5 PCs for the 5 – 15 Hz band, and 7, 27, 84 PCs for the 15 – 30, 30 – 60, 60 – 150 Hz bands, respectively. Velocity was calculated as the Euclidean distance between adjacent time points (at 600 Hz sampling rate) in the corresponding N-dimensional PC space divided by the incremental time unit (Δt = 1.67 ms). Similar to the SCP range, we computed velocity for both trial-averaged and single-trial trajectories.

### Support Vector Machine (SVM) decoding analysis

For each subject, single-trial classification of “seen” vs. “unseen” was performed using activity from all sensors. Using the LIBSVM package [62], we implemented a SVM at each time point around stimulus onset. A five-fold cross-validation scheme was applied, using five interleaved sets of trials. Trials were balanced in the training set by using a random subset of trials in which the number of trials was equalized between the two conditions. Classification was performed 10 times, each time using a different random subset of balanced trials for training, and performance was averaged across iterations. Classification performance for each subject was reported as the average across the five folds. To determine the optimal value of the cost parameter C of the linear decoder, we investigated the cross-validation performance of the decoder for classifying seen vs. unseen trials at all time points for values of C ranging from 2^-10, 2^-8, …, 2^18, following a method recommended by the authors of the LIBSVM toolbox (http://www.csie.ntu.edu.tw/~cjlin/papers/guide/guide.pdf, section 3.2). Seen vs. unseen classification was similar for all values of C but was optimized by setting C = 2^-6. For simplicity and uniformity of analysis, we used C = 2^-6 for all decoding analyses. We additionally assessed the temporal generalization of classification accuracy by training the classifier at each time point and testing on all time points within the epoch. Group-level statistical significance of classifier accuracy was established using cluster-based permutation test. Clusters were defined as contiguous time points where the p-value assessed by a one-tailed Wilcoxon signed rank test against chance level (50%) was less than 0.05. The test statistic W of the Wilcoxon signed rank test was summed across time points in a cluster to yield that cluster’s summary statistic. Null distribution was derived by randomly permuting trial labels for each subject one hundred times. Clusters in the original data were considered significant if their summary statistic exceeded the 95th percentile of the null distribution.

Activation patterns corresponding to the MEG activity contributing to the classifier were computed for each subject and time point by multiplying the vector of SVM decoder weights with the covariance matrix of the data set used to train the classifier [35]. For display purposes, activation patterns were averaged across subjects and scaled separately for each time point.

### Sensor- and PC - Level Analyses on Across-Trial Variability

Across-trial variability of SCP was calculated as s.d. across trials. To visualize changes in variability from baseline, the mean of the pre-stimulus baseline (from -1 sec to -250 ms) was subtracted from each time point. Volume of the state space was calculated as the product of across-trial s.d. across the top 5 PCs [20], and expressed as %change from the baseline. Significant changes were determined using a one-sample t-test against 0 across subjects. Correction for multiple comparisons was carried out using cluster-based permutation tests as described above, except for a two-tailed test. Thus, temporal clusters were defined as contiguous time points that exhibited a significant (*p* < 0.05) difference from 0 and whose t-statistics had the same sign. The absolute value of the summed t-statistic across time points in a cluster was defined as that cluster’s summary statistic. Null distribution was constructed by randomly shuffling the assignment of the post-stimulus and baseline labels 1000 times for each subject. Clusters in the original data were considered significant if their summary statistic exceeded the 2.5 percentile value of the null distribution (corresponding to *p* < 0.05 for a two-tailed test). We also directly compared %change in volume between each pair of task conditions by using paired t-tests, which were corrected for multiple comparisons using cluster-based permutation tests similar to that described above, except that condition labels were randomly shuffled for each subject.

### Across-Trial Variability Based on the Norm or Angle of Population Activity

The location of population activity in the state space can be described by the angle and norm of the vector that points to it. We computed the across-trial variability of the angle and norm of population activity in the 0.05 – 5 Hz frequency band. To compare with the volume analysis, the first five PCs were used. Activity at every time point was then defined by a vector in a 5-dimensional space. The norm of this vector is calculated as

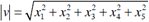

where *x*_*i*_ defines the activity of principal component *i*. The across-trial variability of the norm was calculated as its s.d. across trials. To calculate the across-trial variability of the angle, at each time point we computed the angle (α) between two vectors (*v*^*A*^ and *v*^*B*^) corresponding to the population activity in two different trials as follows:

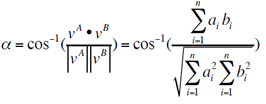

where *a*_*i*_and *b*_*i*_represent the activity from principal component *i* in trial *A* and *B*, respectively. The top five PCs were used (thus, n = 5). The mean α, measured across all trial-pairs within each condition, defined the across-trial variability of the angle.

Separately for seen and unseen trials, we tested whether across-trial angle variability differed between each post-stimulus time point and the pre-stimulus period (from 1 sec before to stimulus onset) by using a non-parametric two-sample test for equal medians [63] as implemented in circ_cmtest function of the CircStats toolbox in MATLAB [64]. Correction for multiple comparisons was carried out using a cluster-based permutation test, where the null distribution was constructed by randomly shuffling the assignment of the post-stimulus and baseline labels. In addition, angle variability was compared between seen and unseen conditions at each time point using the same non-parametric two-sample test for equal medians, and corrected by cluster-based permutation tests based on shuffling condition labels.

### Single-Trial Decoding Based on the Norm or Angle of Population Activity

Norm and angle of population activity in the SCP range (0.05 – 5 Hz) were used. Cross-validation was performed by an interleaved odd-even split. For the training set, the activity of every sensor was averaged across trials within each condition (seen vs. unseen). This yielded a 1 x 273 vector for both seen and unseen conditions. For each trial in the test set, the difference in its angle or norm relative to the “seen” or “unseen” vector from the training set was calculated and the trial was classified as either condition based on the smaller value. We assessed significance of classifier accuracy at each time point using a cluster-based permutation test similar to the one applied to SVM classifier accuracy, as described above.

### The Influence of Pre-Stimulus Activity on Objective Performance in Unseen Trials

Broadband data in the 0.05 – 150 Hz range were used for this analysis (results obtained using SCPs were very similar). First, stimulus-related activity templates were generated for each subject by averaging activity patterns in a 100 ms post-stimulus time window across unseen trials, separately for each stimulus orientation. For each trial, spatial correlation was then computed between the mean MEG activity in a 100 ms pre-stimulus time window and the two templates. Trials were sorted by whether their pre-stimulus activity was more similar to (i.e. had a larger Pearson correlation for) the “left tilt” stimulus template or the “right tilt” stimulus template.

This procedure was performed using all 273 sensors, as well as using sensors from each lobe separately. For the latter analysis, sensors were assigned to lobes according to the sensor naming convention for the CTF MEG scanner, which groups sensors into occipital, temporal, parietal, central, and frontal regions. The number of trials in the two groups were not significantly different (all *p* > 0.5). For each group of trials, we used the classic SDT approach [46] to calculate subjects’ response sensitivity (d’) and bias (c) in their orientation discrimination performance. This analysis was repeated using window lengths of 200, 300, 400 and 500 ms for the post-stimulus template definition and the corresponding pre-stimulus baseline period.

## ACKNOWLEDGEMENTS

This research was supported by the Intramural Research Program of the National Institutes of Health/National Institute of Neurological Disorders and Stroke and New York University Langone Medical Center. We thank Qi Li for sharing the data used in this study, Rishidev Chaudhuri, Richard Hardstone and Ella Podvalny for comments on a previous draft of the manuscript. BJH also acknowledges support by Leon Levy Foundation and Klingenstein-Simons Fellowship.

